# Single-molecule nanopore dielectrophoretic trapping of α-Synuclein with lipid membranes

**DOI:** 10.1101/2022.01.24.477462

**Authors:** Jinming Wu, Tohru Yamashita, Andrew D. Hamilton, Sam Thompson, Jinghui Luo

## Abstract

The lipid-α-Synuclein (α-Syn) interaction plays a crucial role in the pathogenesis of Parkinson’s disease. Here, we trap α-Syn at a conjunction of an α-hemolysin (αHL) single nanopore-lipid to investigate the folding and unfolding kinetics of α-Syn in a lipidic environment. The hybridized α-Syn is generated through a reaction between a 5’-thiol-modified nucleotide oligo (dC30) and the α-Syn mutant (A140C). Owing to an applied voltage, single-molecule hybridized α-Syn can be trapped at the single nanopore. The trapping events are associated with dielectrophoretic force. The folding and unfolding events of α-Syn can be observed at the pore-membrane junction through interpretation of blockade current amplitudes and dwell time. This can be related to the protein quaternary structure influenced by the α-Syn-membrane interaction, allowing further analysis of α-Syn conformational dynamics. We studied how disease associated metal ions (Cu^2+^, Zn^2+^) modulate folding and unfolding of α-Syn at the interface of the membranes and pore, and how α-helical peptidomimetics stabilize the helical conformation of α-Syn in the presence of a membrane. These studies aid our understanding of the complexity of the interaction of α-Syn, lipid membranes and metal ions, and in using peptidomimetics, a new strategy against α-Syn toxicity and aggregation is advanced.

## Introduction

Contributing to the pathogenesis of Parkinson’s disease (PD), intrinsically disordered α-Synuclein (α-Syn), a 140-residue protein, is extensively expressed in neurons and enriched in the synaptic cleft^1^. α-Syn consists of 3 domains: a membrane binding region with positively charged N-terminal residues from 1-60^2,3^, an aggregation associated central hydrophobic domain (NAC: from residues 61-95) and a disordered acidic C-terminal residues from 96-140^4,5^. During aging, the protein deposits as β-sheet rich amyloid fibrils^6^, with contemporaneous neuronal dysfunction and degeneration in the brain of PD. Although the biological role of α-Syn remains elusive, a number of studies suggest that it interacts with phospholipid membranes in physiology and pathology, such as synaptic regulation and neuronal death^7,8^. In an effort to reveal the molecular basis of α-Syn toxicity and aggregation, the interaction of α-Syn with membranes has been widely explored^9–11^. Several models have been presented to explain the α-Syn induced toxicity to lipid membranes: (1) mem-brane-permeabilizing toroidal or barrel pores^12^; (2) carpet model of disrupting and thinning membrane^13,14^; and (3) lipid extraction model^15^. Reciprocally, the nature of the lipid membrane affects the misfolding and aggregation of α-Syn^16^. Folding and unfolding of α-Syn in a lipid membrane environment plays a vital role in toxicity yet the kinetics remain to be fully explored. Metal ions also act as an important factor to modulate the folding, aggregation and toxicity of α-Syn^17^. The cerebrospinal fluid of the substantia nigra from PD patients has abnormally high concentrations of Cu^2+^, Fe^3+^ and Zn^2+18–20^. This suggests that metal ions are involved into the interaction between α-Syn and lipid membranes in the pathogenesis of PD, either in a direct causal manner, or as a consequence of misfolding. Thus, it would be interesting to investigate how metal ions modulate the folding of α-Syn in a lipidic environment.

Single-nanopore technologies have been used to record the interaction or aggregation among amyloid proteins with or without metal ions at the single-molecule level^21–23^. The recording is on the basis of individual amyloid proteins blocking or translocating through a single nanopore, such as an α-hemolysin (αHL) nanopore, in reconstituted lipid membranes across *cis* (ground side) and *trans* sides. Several studies have investigated how amyloid proteins block or translocate the single nanopore for the characterization of amyloid aggregation or interaction^22,23^. The dwell time and residual current of single-nanopore transient blockade by individual amyloid proteins can be extracted to gain an understanding of protein folding, topology and noncovalent interactions of lumen in the nanopore^24,25^. Gurnev *et al*. observed the membrane binding α-Syn of which the C-terminal tail entered the pore and the N and NAC domains partitioned on the surface of the lipid membrane^11^. Tavassoly *et al*. used the same setup and found that Cu^2+^ ions induce large conformational changes of α-Syn^26^. However, in these cases, it is unclear how the whole sequence of α-Syn interacts with the lipid membranes and how metal ions regulate entire α-Syn folding and unfolding on lipid membranes. Recently, Rodriguez-Larrea *et al*. studied the unfolding kinetics of thioredoxin (Trx) in a conjunction with a DNA oligonucleotide leader oligo(dC30) through a nanopore^27,28^. The oligo(dC30) linked Trx was added to the *cis* part of the chamber, where the unfolding of Trx only occurs without interacting with lipid membranes. Their work provides an insight into the unfolding kinetics of Trx without interacting with lipid membranes. Folding and unfolding of α-Syn in a lipid membrane environment plays a vital role in toxicity yet the kinetics remain to be fully explored.

Using single-nanopore analysis, we investigated how Cu^2+^ modulates α-Syn folding and unfolding states on the *trans* side that has a lipidic environment shown in **Fig.1**. By conjuncting α-Syn mutant (A140C) with oligo(dC30), we observed two step-wise blockades of single-nanopore by the oligo-linked α-Syn, which may be explained by a single-nanopore dielectrophoresis (DEP) force model^29^. In this model, two blockade levels represent α-Syn quaternary folding and unfolding structures, caused by its interaction with the lipid membrane and trapped by a strong DEP force at the pore-membrane junction. We further studied two metal ions, Cu^2+^, Zn^2+^ and a helix mimetic compound that modulates α-Syn dynamics in a lipidic environment. These studies seek insights into the complexity of α-Syn interactions with lipid membranes in the presence of metal ions and small molecule modulators of misfolding, thus allowing the development of new strategies against α-Syn toxicity and aggregation.

**Figure 1.**
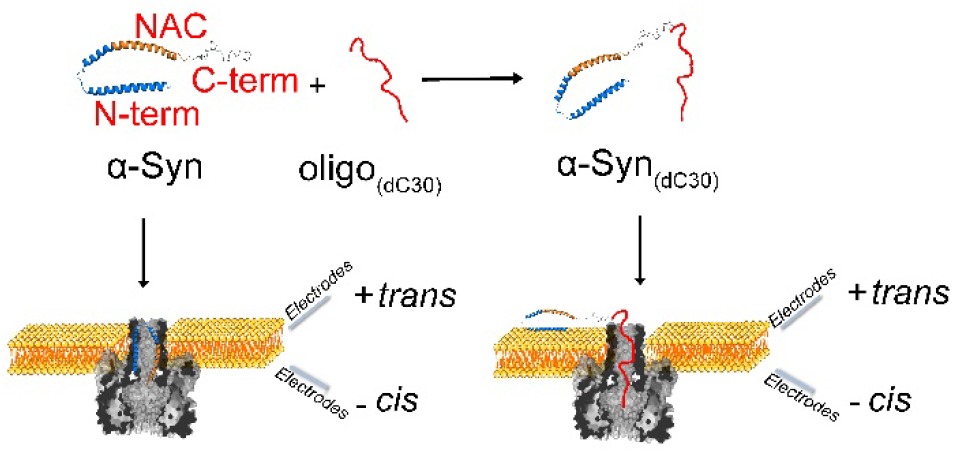
Illustration of α-Syn-membrane interactions studied by single-molecule nanopore analysis. WT α-Syn was added into the *trans* side of a single α-hemolysin (αHL) nanopore and induced the blockage of αHL channel in a lipid bilayer. To study the interaction of full length α-Syn with the lipid membrane, α-Syn mutant (A140C) was linked to oligonucleotides oligo(dC30). The conjuncted α-Syn_(dC30)_ was then added into the *trans* side of αHL nanopore and induced the dielectrophoresis trapping of α-Syn_(dC30)_ in lipid membranes.

## Results and discussion

To investigate whether PD associated Cu^2+^ ions modulate the binding between the lipid bilayer and α-Syn, we conducted single-molecule αHL nanopore electrical recording in the presence of α-Syn with or without Cu^2+^ ions, shown in **Fig. 1**. The αHL nanopore was reconstituted in a planar lipid bilayer, composed of DOPC and DOPG (in a ratio of 4:1) and recorded at −100 mV. **Fig. 2A** shows transient blockade events of the α-Syn without and with Cu^2+^ from the *trans* side, which is consistent with the previous observation that α-Syn causes transient nanopore blockage^23^. In the presence of Cu^2+^, α-Syn induces a lower residue current, which is shown as the red scatter distribution in **Fig. 2B**. The histogram analysis of the residue current amplitudes estimates α-Syn with and without Cu^2+^ ions to be −21 pA and −11.5 pA respectively. The lower nanopore blockage induced by the addition of Cu^2+^ ions revealed a more folded structure of α-Syn. The transient nanopore blockage may be caused by the translocation of α-Syn to the *cis* side^11^. The charged C-terminus of α-Syn presumably leads the translocation of the full-length protein into the lumen of the nanopore. When the C-terminus enters the lumen, the current changes can be mainly attributed to the bilayer interaction with the N-terminus rather than the full-length protein. However, the transient current signal does not provide informative biophysical characterization for single-molecule α-Syn interaction with lipid membrane.

**Figure 2.**
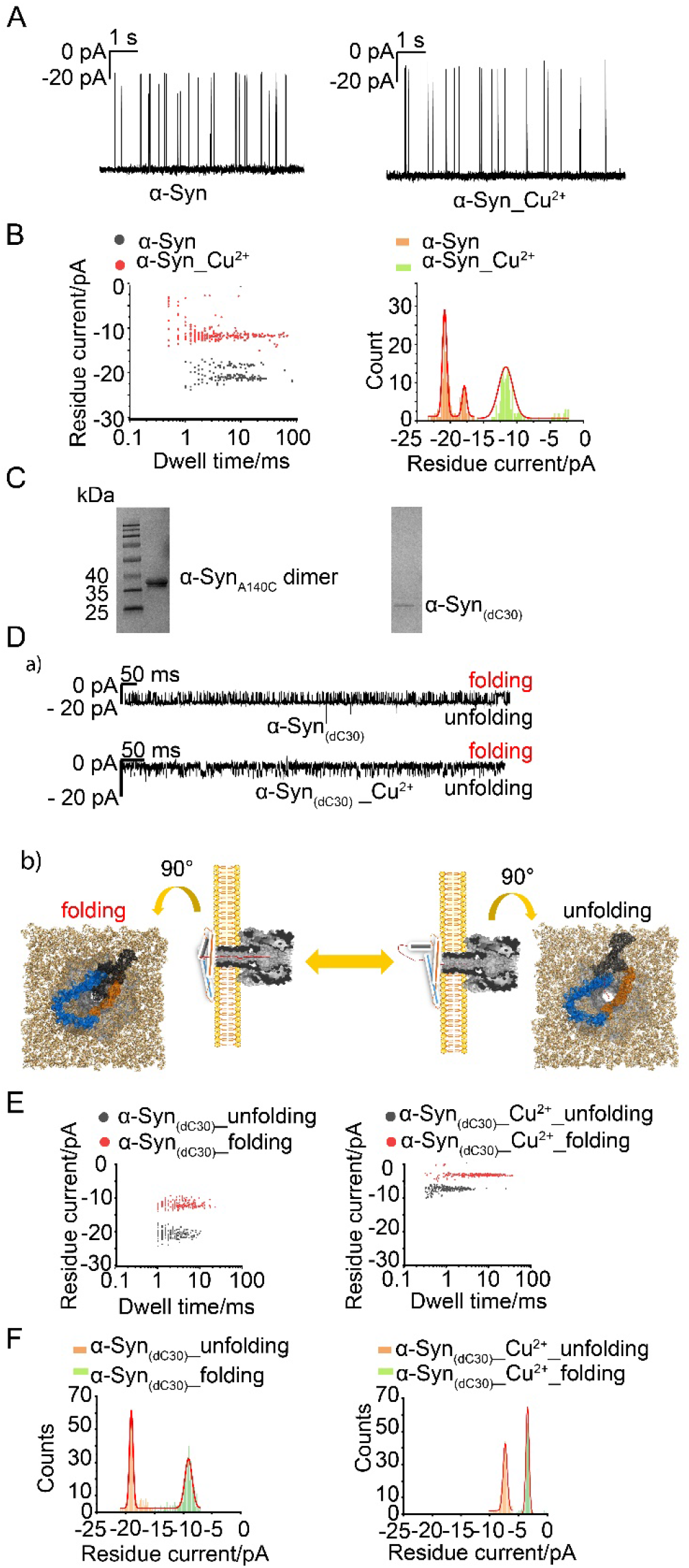
(A) Representative current traces reveal the translocation of wild-type (WT) α-Syn through a single αHL nanopore in the presence or absence of Cu^2+^ ions. The samples were added to the trans side of the lipid bilayer, composed of DOPC and DOPG (4:1). (B) Left: scatter distribution of the event dwell time plotted against the residue current in the presence of WT α-Syn with or without Cu^2+^ ions. Right: residue current histogram in the presence of WT α-Syn with or without Cu^2+^ ions. (C) The conjuncted α-Syn (A140C) with single-strand DNA oligos (dC30) through pyridyl disulfide reaction (Left) and α-Syn (A140C) dimer. (D) a) Representative current recording of single αHL nanopore across a planar lipid bilayer composed of DOPC and DOPG (4:1) in the presence of α-Syn_(dC30)_ with or without Cu^2+^, at −100 mV. b) Illustraion of α-Syn folding and unfolding in lipid membranes. The more stably folded the structure of the α-Syn moiety, the lower the residue current. (E) Scatter distribution of the event dwell time plotted against the residue current in the presence of α-Syn_(dC30)_ with or without Cu^2+^ ions. Due to the nanopore dielectrophoresis trapping, the folded α-Syn_(dC30)_ generated smaller residual current than the unfolded structure. Two distinct events were classified, the folding (red scatters) and unfolding events (black scatters) of α-Syn_(dC30)_. (F) Residue current histogram in the presence of α-Syn_(dC30)_ with or without Cu^2+^ ions. Similar to the presence of WT α-Syn, the addition of Cu^2+^ ions to the hybrid α-Syn reduced the residue current of electrical recording, suggesting the different α-Syn conformations in the presence of Cu^2+^ ions.

To gain insights into the interaction of full-length α-Syn with lipid membranes, we linked α-Syn (A140C) to a single-strand DNA oligomer (dC30) using a pyridyl disulfide reaction^28^. As shown on SDS-PAGE in **Fig. 2C**, we purified the hybrid α-Syn_(dC30)_, which was further confirmed by LC-MS (**Fig. S3**). The addition of α-Syn_(dC30)_ to the *trans* side caused the trapping of αHL nanopore in **Fig. 2D**. This may be attributed by two factors: (1) α-Syn_(dC30)_ released the full-length α-Syn for the interaction with the head group of lipid membranes; (2) with its longer sequence, α-Syn_(dC30)_ had a stronger dielectrophoretic force (DEP) at the nanopore conjunction than the WT. DEP trapping was observed in a previous study where α-Syn could be trapped in the constriction of a nanopore, and was considered as a reservoir-microchannel junction^23^. Here, we observed two distinguished trapping levels in **Fig. 2D-F**, with approximately 10 pA current difference in the presence of α-Syn_(dC30)_ but not WT α-Syn. At a nanopore conjunction, a DEP of the opposite electric field induces particle deflection, focusing and trapping^29^. Large complexes, like α-Syn_(dC30)_, can be trapped in the conjunction but the smaller WT α-Syn translocate through the nanopore. Two different trapping events can be induced by the α-Syn interaction with the lipid surface and a DEP that contributed to the folding and unfolding of α-Syn on the lipid membranes. The lower the residue current we observed, the more stably folded the structure of the α-Syn moiety is anticipated to be. Different residue current levels suggest that α-Syn may display dynamics and kinetics with two folding states while trapped at the lipid and nanopore conjunction. The dynamics and kinetics could be modulated by metal ions or small molecule ligands. For instance, shown in **Fig. 2E-F**, Cu^2+^ ions refolded α-Syn in a more compacted structure and reduced residue current in the electrical recording. Analysis of these trapping events offers insights into the conformational dynamics and kinetics of α-Syn in the presence of lipid membranes, metal ions and aggregation inhibitors.

As a control, we investigated the conformational dynamics of α-Syn_(dC30)_ in the presence of neutrally charged lipid membrane, composed of DPhPC. **Fig. 3A** shows that α-Syn displays fewer folding and unfolding events in DPhPC lipids than that in negatively charged DOPC:DOPG (4:1) lipid membranes (**Fig. 2D**), suggesting a less α-Syn folded structure that completely traps the nanopore. This may be explained by greater folding of α-Syn with negatively charged lipid membrane, in agreement with previous studies^30–32^. Similar to the presence of negatively charged lipid membrane, the addition of Cu^2+^ ions and the hybridized α-Syn to the neutrally charged lipid membrane gave reduced residue current in the electrical recording. This reveals both Cu^2+^ ions and the nature of the lipid determine the conformation of α-Syn.

**Figure 3.**
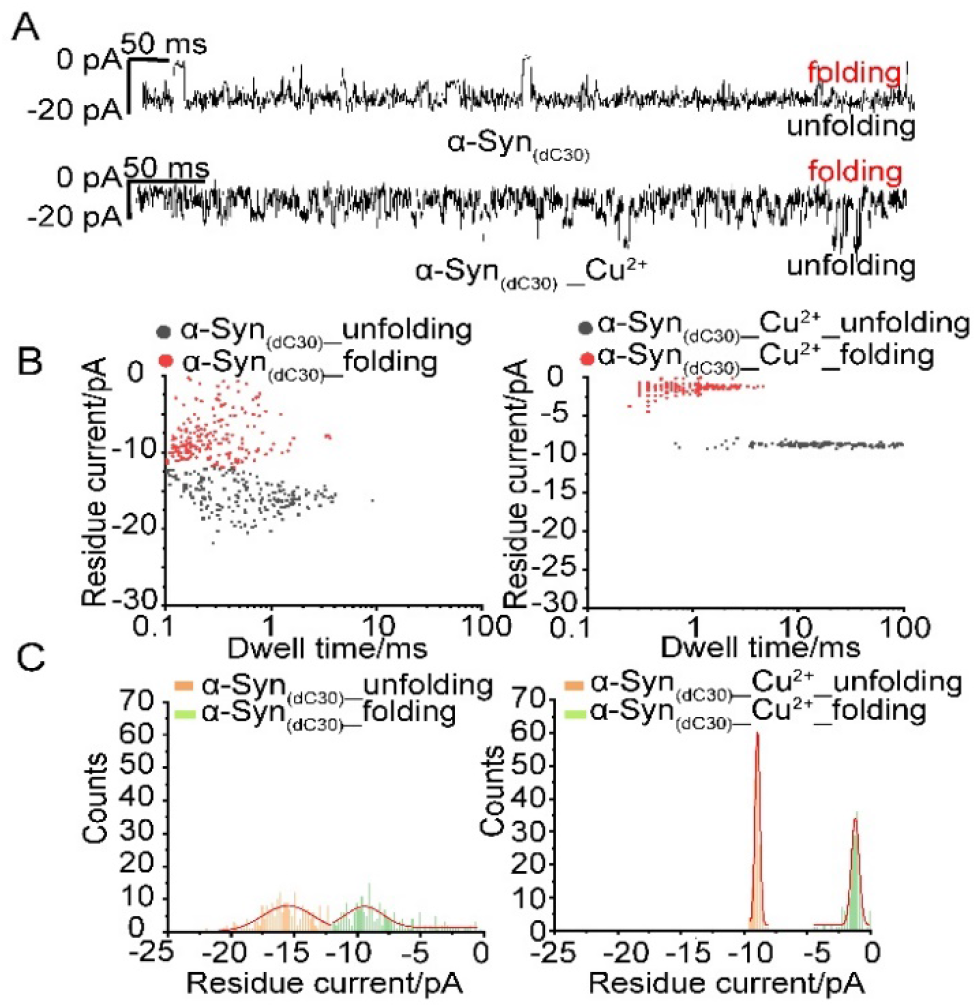
(A) Representative current recording of hybridized α-Syn_(dC30)_ with or without Cu^2+^ translocating a single αHL nanopore on the neutrally charged DPhPC lipid bilayer at −100 mV. Two distinct folded and unfolded events of α-Syn_(dC30)_ were classified. The folded α-Syn_(dC30)_ induced smaller residual current than the unfolded structure. The final concentration of hybridized α-Syn_(dC30)_ and Cu^2+^ is 0.2 μM and 5 μM respectively. (B) Scatter distribution of the dwell time plotted against the residue current of α-Syn_(dC30)_ with or without Cu^2+^. The red and black scatters represent the folding and unfolding events respectively. (C) The distributions of residue current of α-Syn_(dC30)_ with or without Cu^2+^ plotted as histograms, fitting with multiple-peak Gaussian function.

An increased dielectrophoretic force was carried out for studying different trapping events. We increased the voltage to −200 mV and observed in **Fig. 4A** that the trapping events are similar to the recording at −100 mV in **Fig. 2D**. However, the presence of Cu^2+^ ions slightly changes the effect of trapping at −200 mV in **Fig. 4A**, in comparison to the trapping at −100 mV. It suggests a higher voltage may give a stronger dielectrophoretic force for the trapping and interaction of the α-Syn moiety with the lipid membranes in the presence of Cu^2+^ ions. A plausible explanation is that α-Syn forms a folded complex with Cu^2+^ ions, taking a stronger dielectrophoretic force at a higher voltage. We compared the modulation of the α-Syn interaction with lipid membranes in the presence of Cu^2+^ and Zn^2+^ at −200 mV. Since the lowest blockade current is observed with Cu^2+^ this suggests a more compacted α-Syn conformation. Likewise, the folding time constant is higher with Cu^2+^ than Zn^2+^ (**Table 2**), suggesting Cu^2+^ forms a more stable complex with α-Syn in the lipid membrane. In our trapping experiment, the residue current and time constant provide insight into the folding and complex stability information of metal ions, α-Synuclein and lipid membranes.

**Figure 4.**
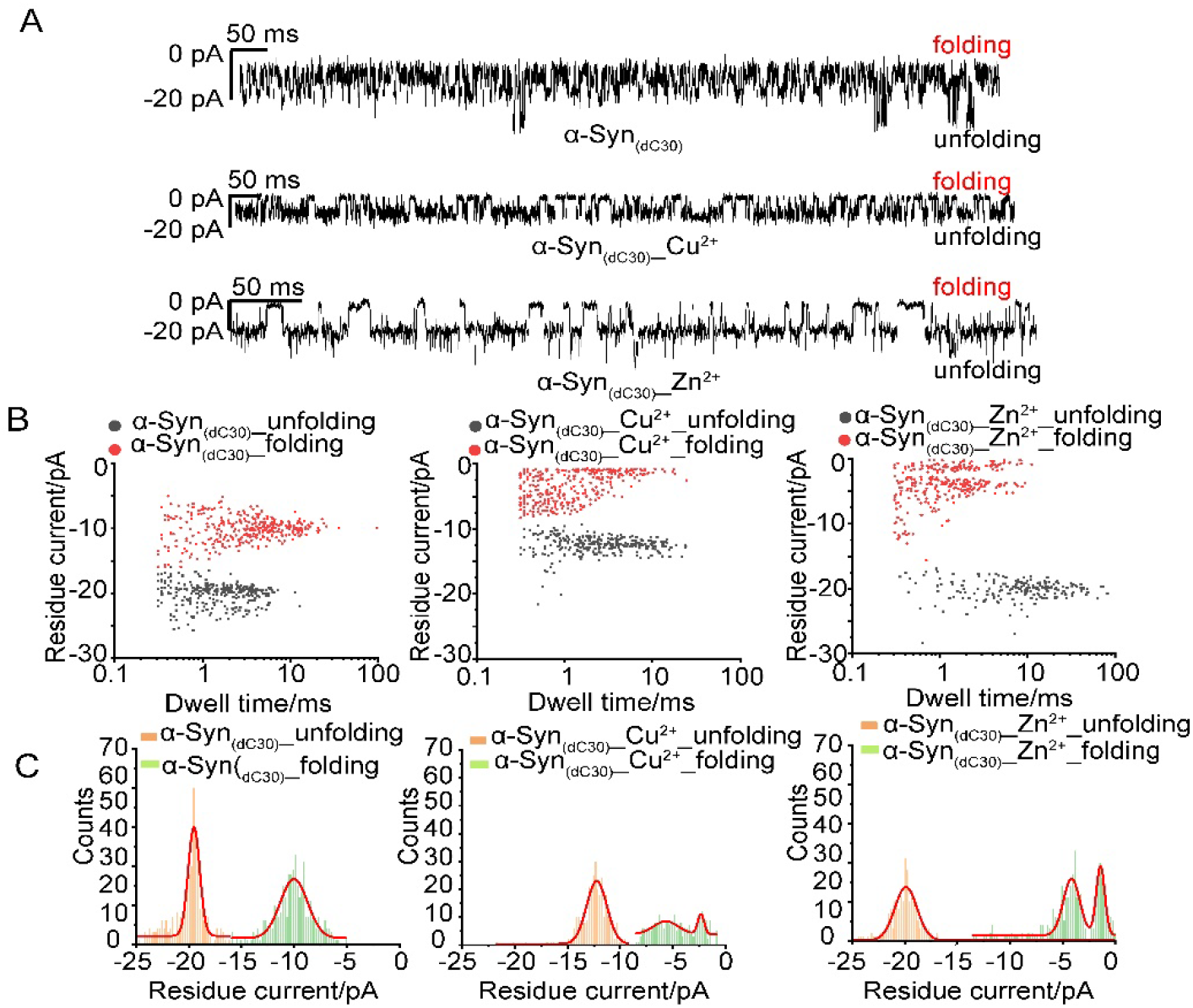
(A) Representative current recording of hybridized α-Syn_(dC30)_ with or without Cu^2+^ or Zn^2+^ translocating a single αHL nanopore on the negatively charged DOPC:DOPG (4:1) lipid bilayer at −200 mV. Two distinct folded and unfolded events of α-Syn_(dC30)_ were classified. The folded α-Syn_(dC30)_ induced less blocked current than the unfolded structure. The final concentration of hybridized α-Syn_(dC30)_ and Cu^2+^ or Zn^2+^ is 0.2 μM and 5 μM respectively. (B) Scatter distribution of the dwell time plotted against the residue current of α-Syn_(dC30)_ with or without Cu^2+^ or Zn^2+^. The red and black scatters represent the folding and unfolding events respectively. (C) The distributions of residue current of α-Syn_(dC30)_ with or without Cu^2+^ or Zn^2+^ plotted as histograms, fitting with multiple-peak Gaussian function.

**Table 1.**
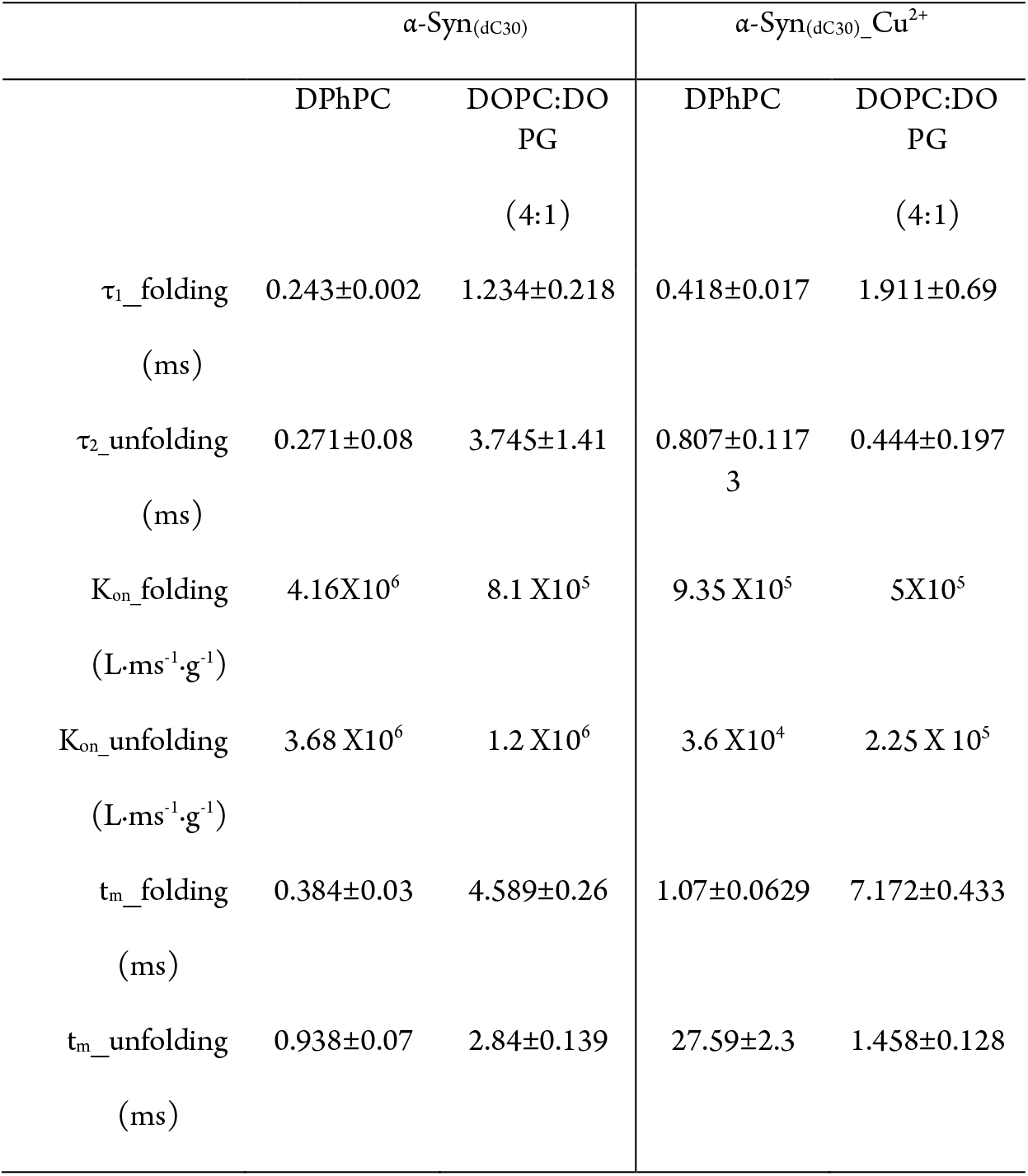
The folding and unfolding events of α-Syn_(dC30)_ with or without Cu^2+^ at −100 mV on DPhPC and DOPC:DOPG lipid bilayer. τ1_folding and τ2_unfolding are calculated from the scatter distribution of the dwell time against the residue current, which were fitted by the double-exponential decay function. The rate constant of K_on_=1[τ_on_×C_peptide_]. t_m_ represents the mean value of the dwell time.

**Table 2.**
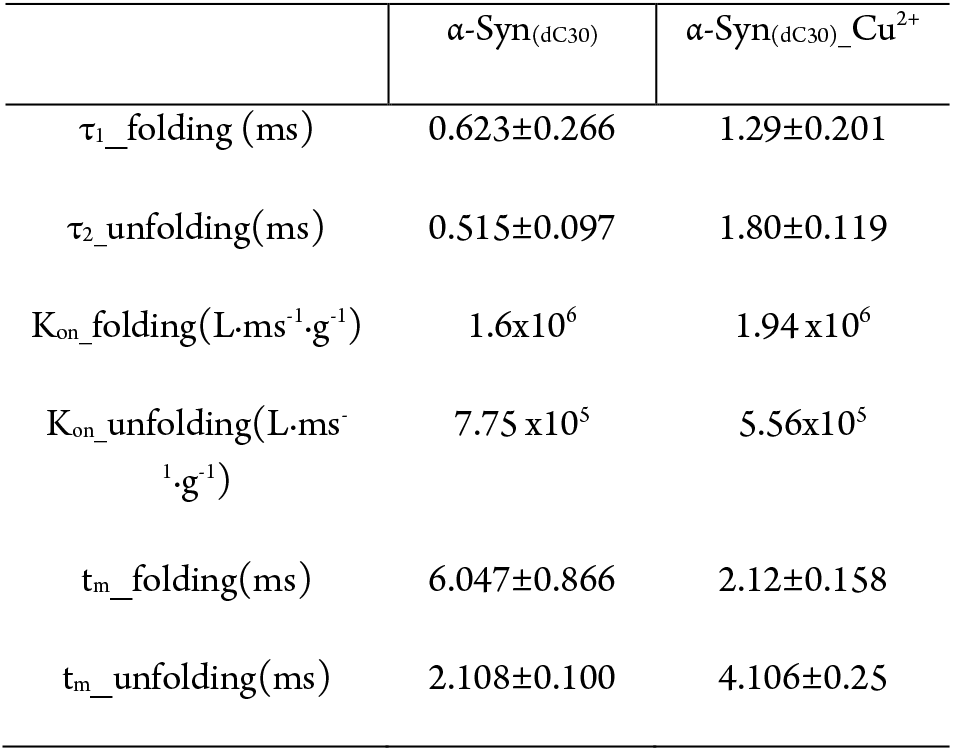
The folding and unfolding events of α-Syn_(dC30)_ with or without Cu^2+^ at −200 mV on DOPC:DOPG (4:1) lipid bilayer

| | $\alpha\text{-Syn}_{(\text{dC30})}$ | $\alpha\text{-Syn}_{(\text{dC30})}\text{-Cu}^{2+}$ |
| --- | --- | --- |
| $\tau_{\text{folding}}(\text{ms})$ | $0.623 \pm 0.266$ | $1.29 \pm 0.201$ |
| $\tau_{\text{unfolding}}(\text{ms})$ | $0.515 \pm 0.097$ | $1.80 \pm 0.119$ |
| $K_{\text{on\_folding}}(\text{L}\cdot\text{ms}^{-1}\cdot\text{g}^{-1})$ | $1.6 \times 10^6$ | $1.94 \times 10^6$ |
| $K_{\text{on\_unfolding}}(\text{L}\cdot\text{ms}^{-1}\cdot\text{g}^{-1})$ | $7.75 \times 10^5$ | $5.56 \times 10^5$ |
| $t_{\text{m\_folding}}(\text{ms})$ | $6.047 \pm 0.866$ | $2.12 \pm 0.158$ |
| $t_{\text{m\_unfolding}}(\text{ms})$ | $2.108 \pm 0.100$ | $4.106 \pm 0.25$ |

The single-nanopore trapping technique was further applied to investigate how small molecules, which can disrupt amyloid protein fibrillization kinetics, modulate α-Syn conformation in the presence of a lipid membrane (**Fig. 5**). Compound **3**, an α-helical mimetic compound that can imitate the topography of an α-helix, is functionalized with, at physiological pH, cationic NH_3_^4^, anionic COO^-^, and branched alkyl groups in the *i*, *i*+*4* and *i*+*7* positions respectively (**Fig. 5B**)^33^. Mimetic **3** is designed to target the helical surface of α-Syn comprising three amino acids (negatively charged Glu^46^, positively charged His^50^ and hydrophobic Ala^53^) that occupy the *i*, *i* + *4* and *i* + *7* positions (**Fig. 5A**), by forming complementary contacts. We have previously successfully demonstrated such an approach with islet amyloid polypeptide^34^. As shown in **Fig.5C**, compared to α-Syn_(dC30)_ alone, the folding and unfolding events of α-Syn in the presence of mimetic **3** in lipid membranes is significantly reduced, consistent with the stabilization of helical protein. Besides peptidomimetic **3**, dopamine, another compound that kinetically stabilizes α-Syn oligomers^35–38^, also reduces the folding and unfolding events of α-Syn in lipid membranes, but with more spikes and greater residual current, indicative of the weaker binding of dopamine to α-Syn than that of mimetic **3**. These results suggests singlenanopore trapping is a suitable method to characterize the effect small molecules exert on α-Syn conformational dynamics in the presence of lipid membranes. Moreover, it may find use as a primary tool for the screening and optimization of aggregation inhibitors.

**Fig. 5.**
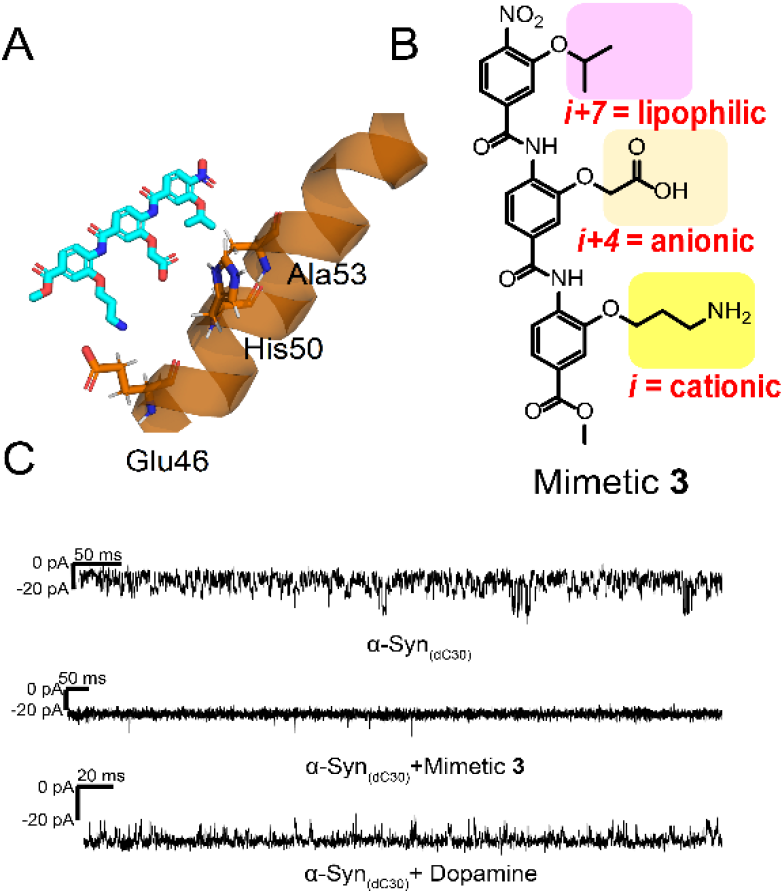
(A) Schematic representation of an α-helix mimetic compound interacting with three residues (Glu^46^, His^50^, and Ala^53^) located in the α-helical region of membrane-bound α-Syn. (B) Structure of α-helix mimetic compound **3** with three side chains (yellow, orange and pink highlights) designed to form complementary contacts with residues Glu^46^, His^50^ and Ala^53^ through salt bridges at *i* and *i*+*4*, and hydrophobic interactions at *i+7*. (C) Representative current recordings of hybridized α-Syn_(dC30)_ in the presence of mimetic **3** or dopamine. Negatively charged DOPC:DOPG (4:1) lipid bilayer was formed under the voltage of −200 mV.

## Conclusions

Previously, single-nanopore studies characterized the conformation and dynamics of amyloid proteins using a number of heterogeneous events rather than the kinetics of a single-molecule amyloid protein. Here we observed the α-Syn interaction with lipids at a single-molecule level by trapping the molecule under externally applied dielectriphoretic force. We show that the folding and unfolding kinetics of α-Syn in the presence of lipid membranes can be extracted from single-molecule nanopore dielectrophoretic trapping. The folding and unfolding of α-Syn is modulated by the charged nature of the lipids constituting the membranes and the metal ions. Therefore, the nanopore trapping method offers a single-molecule detection for understanding the folding and unfolding conformation of intrinsically disordered proteins in a lipidic environment.

The limitations of our study are that folding and unfolding characteristics of trapping are for hybridized α-Syn under externally applied dielectriphoretic force. The external force and hybridization may influence the intrinsic structural properties of α-Syn in a lipidic environment. To address these issues future work in our laboratory will explore the use of electrode-free nanopore sensing^39^.

Last, we found that disease associated metal ions and peptidomimetic compounds modulate the folding dynamics of α-Syn in lipid membranes. In principle, the method can be used for screening inhibitors suppressing the interaction between lipid and α-Syn, which will be the subject of further study. We believe that further parallelization of single-molecule nanopore trapping has the potential to enable high-throughput screening of potential therapeutics against intrinsically disordered proteins in lipidic environments.

## ASSOCIATED CONTENT

### Supporting Information

The experimental section and Figures S1-9 are available in supporting information.

## ACKNOWLEDGMENT

We acknowledge financial support from The Universities of Oxford and Southampton, the Paul Scherrer Insitute, the EPSRC (EP/S028722/1, S.T.), the Swiss National Scientific Foundation ((310030_197626, J.L.), the Brightfocus foundation (A20201759S, J.L.) and Takeda Pharmaceutical Company Limited for their generosity in providing financial and logistical support during a sabbatical position for T.Y. as a visiting Scientist in The University of Oxford.

## Notes

### Competing Interest Statement

The authors have declared no competing interest.

