## Supplemental information for "Single-molecule nanopore dielectrophoretic trapping of α-Synuclein with lipid membranes"

**Abstract: The lipid-α-Synuclein (α-Syn) interaction plays a crucial role in the pathogenesis of Parkinson’s disease. Here, we trap α-Syn at a conjunction of an α-hemolysin (αHL) single nanopore-lipid to investigate the folding and unfolding kinetics of α-Syn in a lipidic environment. The hybridized α-Syn is generated through a reaction between a 5’-thiol-modified nucleotide oligo (dC30) and the α-Syn mutant (A140C). Owing to an applied voltage, single-molecule hybridized α-Syn can be trapped at the single nanopore. The trapping events are associated with dielectrophoretic force. The folding and unfolding events of α-Syn can be observed at the pore-membrane junction through interpretation of blockade current amplitudes and dwell time. This can be related to the protein quaternary structure influenced by the α-Syn-membrane interaction, allowing further analysis of α-Syn conformational dynamics. We studied how disease associated metal ions (Cu2+, Zn2+) modulate folding and unfolding of α-Syn at the interface of the membranes and pore, and how α-helical peptidomimetics stabilize the helical conformation of α-Syn in the presence of a membrane. These studies aid our understanding of the complexity of the interaction of α-Syn, lipid membranes and metal ions, and in using peptidomimetics, a new strategy against α-Syn toxicity and aggregation is advanced.**

Experimental Procedures

Protein expression of wild-type (WT) αHL and WT α-Syn

WT αHL was expressed based on the previous reports ^[1]^. In brief, αHL plasmid was transformed into competent E.*coil* BL21(DE3)pLysS cells. A single colony was cultured in 100 mL LB medium with 100 μg/mL ampicillin at 37 ℃, 180 rpm. Once the E.*coli* reached an OD600 of 0.6, IPTG was added up to a concentration of 1 mM and the culture was incubated at 18 ℃, 180 rpm overnight. The cell pellet was further lysed in 50 mM Tris (pH 8.0), 500 mM NaCl, 10 mM imidazole and 3.8 mM DDM (n-Dodecyl β-D-maltoside, Cat.ID D310S, Anatrace (Ohio, USA) ) on ice. For His-tagged αHL purification, immobilized metal-affinity chromatography (IMAC) was used in a high-affinity TALON (Cobalt) resin (Sigma-Aldrich). The imidazole gradient elution with concentrations of 20 mM, 50 mM and 500 mM was applied for purifying αHL monomers from heptamers in 50 mM Tris buffer (pH 8.0), with 500 mM NaCl and 3.8 mM DDM. The purified proteins were aliquoted and stored at -80 °C. SDS-PAGE gel electrophoresis (Bio-rad, 4-20 % Mini-PROTEAN® TGX™ Precast Protein Gels) at 200 mV was implemented to verify the protein purification. The protein concentration was determined by Nanodrop at 280 nm.

The primary procedure of WT α-Syn plasmid transformation is similar with αHL. IPTG (final concentration 1 mM) was added into the overnight culture of transformed BL21(DE3)pLysS, after the E.*coli* reached an OD600 of 1.04. After 4 h incubation at 37 °C, 180 rpm, the cell pellet was harvested at 4°C. Osmotic shock purification was carried out. 1L cell pellet was suspended in 100 ml, 40% sucrose osmotic shock buffer with 30 mM Tris, pH 7.2, and 2 mM EDTA disodium. After the incubation for 10 min at room temperature, the centrifugation at 12,000 rpm for 20 min was applied for recollecting the pellet, then subjected to 90 ml cold water with 37.5 μl of saturated MgCl2. After 3 min incubation on ice, the periplasm α-Syn proteins were collected in the supernatant by centrifugation at 12,000 rpm for 20 min. After dialyzed in 20 mM Tris buffer, pH 8 for overnight, α-Syn was eluted in a 0-0.5 M NaCl gradient in 20 mM Tris, pH 8.0 by ion exchange chromatography (IEX) on a 5 mL HiTrap Q-Sepharose Fast Flow column (GE Healthcare). Fractions containing large amount of α-Syn were collected and precipitated by adding the saturated ammonium sulfate solution (4.3 M at room temperature) until 50% saturation (1:1) was reached. The precipitated α-Syn was dissolved in 20 mM Tris (pH 7.2) and subjected to size exclusion chromatography (SEC) on a GF Hiload Superdex 75 16/60 column (GE Healthcare), and eluted with 20 mM Tris, pH7.2. After the SDS-PAGE gel electrophoresis (4–20% Mini-PROTEAN® TGX™ Precast Protein Gels), fractions containing α-Syn were aliquoted and stored at -80 °C.

The purification of α-Syn conjugated with nucleotide oligo (dC30)

The residue Ala^140^ of α-Syn was mutated to Cysteine by using a Quick Change II XL site-directed mutagenesis kit (Stratagene) and confirmed by the sequencing. The plasmid containing α-Syn A140C was expressed and purified as WT α-Syn protein production mentioned above. Prior to the conjunction, α-Syn A140C protein was reduced in the presence of DDT and purified by using size exclusion chromatography (Superdex 75 Increase 10/300 GL, GE Healthcare). Oligo(dC)30 was reacted with α-Syn A140C by using established protocol reported previously ^[2]^. Briefly, 2,2'-Dipyridyldisulfide (Sigma-Cat.ID: 8411090005) was used to activate 5’-thiol (hexamethylene linker)-modified oligo(dC)30 (Integrated DNA Technologies) and 10 mM DTT was subsequently applied for the reduction of the activated oligos for 1 hour. The oligos were separated from DTT by size-exclusion chromatography (Superdex 75 Increase 10/300 GL, GE Healthcare) and reacted with purified α-Syn A140C for 16 h at room temperature. The final product was purified by 0-1 M KCl gradient elution with HiTrap Q FF ion exchange column, GE Healthcare) in 10 mM Tris pH 8.0 with 1 mM EDTA and verified by SDS-Page gel and LC-MS.

LC-MS (Liquid chromatography–mass spectrometry)

The molecule weight of purified proteins including WT αHL, α-Syn and nucleotide oligo (dC30) conjuncted α-Syn were identified by LC-MS. ESI-TOF MS (LCT Premier Mass Spectrometer, Waters AG, Baden-Dättwil, Switzerland) was combined with the LC (Waters 2795). A gradient of ACN/water in the presence of 0.1% formic acid was prepared for the MassPREP Phenyl Guard Column (Waters n°186002785) or the C18 Aeris widepore column (Phenomenex). The obtained MS spectrums for multiply charged protein ions were deconvoluted by using MAxent1 software to obtain the protein mass. The LC-MS results were shown in the supporting information Fig.S3.

Preparation of lipid bilayer

A mixture of 20 mg/mL DOPC (1,2-dioleoyl-sn-glycero-3-phosphocholine, Cat.ID 850375P, Avanti Polar Lipids (Alabaster, AL)):DOPG (1,2-dioleoyl-sn-glycero-3-phospho-(1′-rac-glycerol), Cat.ID 840475C, Avanti Polar Lipids (Alabaster, AL)) (in a ratio of 4:1) in chloroform was prepared in a glass vial, and dried by the N_2_ gas. To completely remove the chloroform, the glass vial was transferred to a vacuum desiccator for 4 h. Pentane (99%, Sigma-Aldrich,) was added to dissolve the dried lipid film of DOPC:DOPG mixture or DPhPC lipid powder at a concentration of 10 mg/mL. The black lipid bilayer was formed by using Montal and Mueller method ^[3]^ with 5 µL DOPC:DOPG (4:1) or DPhPC in two compartments (*cis* or *trans*) of the home-made chamber. The chamber was separated by a 25 μm thick Teflon film (Good Fellow Inc., #FP301200) with a 100 µm aperture. The single-channel electrical recording was carried out in 500 µL, 10 mM Hepes buffer, pH 7.4, with 1 mM KCl. The aperture was pretreated with 0.5 µL, 2 % (v/v) hexadecane/pentane mixture and was dried immediately by N_2_. The electrical signals were collected through a pair of Ag/AgCl electrodes separately into the *cis*- and *trans*-compartments. The *cis* side was defined as the grounded side and a voltage potential was applied to the *trans* side. It means that the positively charged analytes can translocate from *trans* to *cis* through the bilayer.

Single-channel recording and data analysis

The	single-channel recordings were conducted in a whole cell mode with a patch clamp amplifier (Axopatch 200 B, Axon instrument, Molecular Devices, CA). For data acquisition, a DigiData 1440 A/D converter (Axon) was equipped with a PC, where the pClamp and Clampfit were installed for data process. The purified αHL in 50 mM Tris buffer, pH 8.0, with 500 mM NaCl, 500 mM imidazole and 3.8 mM DDM, was diluted to 1 μg/mL. 2 μL αHL proteins were added to the *cis* part of the chamber for a single nanopore formation. The purified protein was added into the *trans* part of the chamber with the final concentration 0.2 μM. The ratio of protein and metal ions is 1:25. All the experiments were carried out at room temperature.

Synthesis of helix mimic compound 3

Reactions were carried out under a nitrogen or argon atmosphere in oven-dried glassware unless otherwise stated. Standard inert atmosphere techniques were used in handling all air and moisture sensitive reagents. Tetrahydrofuran (THF), dichloromethane (DCM), N,N’‑dimethylformamide (DMF) and methanol (MeOH) were anhydrous (dried on an MB-SPS-800 solvent purification system). Other solvents and reagents were used directly as received from commercial suppliers. All aqueous solutions were saturated unless specified otherwise.

Flash column chromatography was carried out using Merck 60 silica gel. Thin-layer chromatography was carried out using Merck Kieselgel 60 F254 (230-400 mesh) fluorescent treated silica, visualized under UV light (254 nm) or by staining with aqueous potassium permanganate solution.

^1^H and ^13^C NMR spectra were recorded using a Bruker spectrometer (400, 500 or 600 MHz) running TopSpin™ software and are quoted in ppm for measurement against residual solvent peaks. Chemical shifts (δ) are given in parts per million (ppm) and coupling constants (*J*) are given in Hertz (Hz). The ^1^H NMR spectra are reported as follows: δ (number of protons, multiplicity, coupling constant). Multiplicity is abbreviated as follows: s = singlet, d = doublet, t = triplet, q = quartet, quint. = quintet, m = multiplet, br = broad. IR spectra were recorded on a Bruker Tensor 27 FT-IR spectrometer from a thin film deposited onto a diamond ATR module. Only selected maximum absorbances (ν_max_) of the most intense peaks are reported (cm^-1^). High-resolution mass spectra were using a Bruker MicroTof (ESI) or an Agilent 7200 Accurate Mass Q-TOF GC/MS with an MSD Direct Inlet Probe (ammonia CI). Compound names are those generated by ChemBioDraw™ (CambridgeSoft) following IUPAC nomenclature.

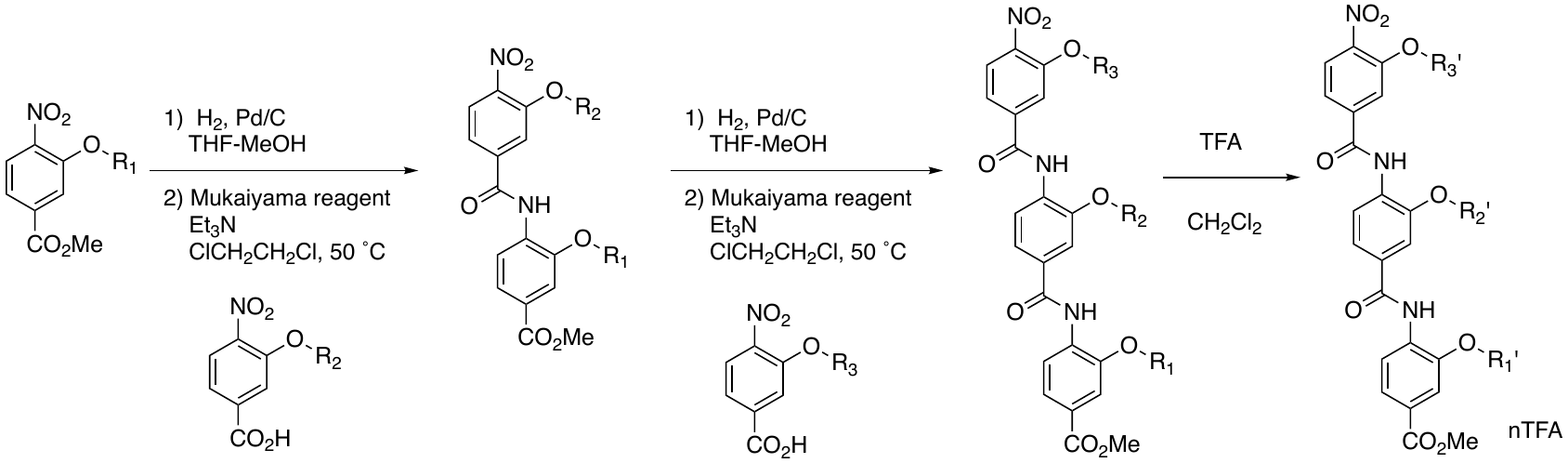
1 General procedure for synthesis of trimeric benzamides

2 General procedure (a) – reduction of nitro group followed by condensation with acid

A mixture of nitro aromatic (1.0 eq) and palladium on carbon (10 % Pd by weight, 50 weight %) in THF-methanol (1:1, 0.4 M) was stirred under a hydrogen atmosphere (balloon pressure) for 2 h. Filtration over Celite™ and concentration in vacuo gave aniline, which was used in the next reaction without further purification.

To a solution of carboxylic acid (1.2 eq) and Et_3_N (2.4 eq) in 1,2-dichloroethane (0.5 M) was added Mukaiyama reagent (2-chloro-1-methylpyridinium iodide, 1.2 eq). The mixture was stirred at 50 ˚C for 15 min. Then a solution of the corresponding aniline (1.0 eq) in 1,2‑dichloroethane (0.5 M) was added to the reaction mixture and the resulting solution was stirred at 50 ˚C for 14 h. The reaction was concentrated *in vacuo* and the residue was purified by flash column chromatography (EtOAc/hexane).

3 General procedure (a’) – reduction of nitro group with zinc followed by condensation with acid

To a mixture of nitro aromatic (1.0 eq) and zinc(0) dust (10 eq) in dichloromethane (0.4 M) was added acetic acid (one quarter the volume of dichloromethane) and the mixture was stirred at 40 ˚C for 1 h. The mixture was neutralized with aq. Sodium hydrogencarbonate and extracted with dichloromethane twice. The organic layers were used in the next reaction without further purification.

To a solution of acid (1.2 eq) and triethylamine (2.4 eq) in dichloroethane (0.5 M) was added Mukaiyama reagent (2-chloro-1-methylpyridinium iodide, 1.2 eq). The mixture was stirred at 50 ˚C for 15 min. Then a solution of the corresponding aniline (1.0 eq) in dichloroethane (0.5 M) was added to the reaction mixture and the resulting solution was stirred at 50 ˚C for 14 h. The reaction was concentrated *in vacuo* and the residue was purified by flash column chromatography (EtOAc/hexane).

4 General procedure (b) – deprotection of *t*-butyl group (*N*-Boc and/or *t*-butyl ester)

To a solution of trimeric benzamide in dichloromethane (0.5 M) was added TFA (same volume as dichloromethane) and the mixture was stirred for 6 h. The reaction was concentrated in vacuo and the crude solid was collected and washed with diethyl ether.

5 Synthetic route to 2-(5-((2-(3-aminopropoxy)-4-(methoxycarbonyl)phenyl)carbamoyl)-2-(3-isopropoxy-4-nitrobenzamido)phenoxy)acetic acid 2,2,2-trifluoroacetic acid salt (3)

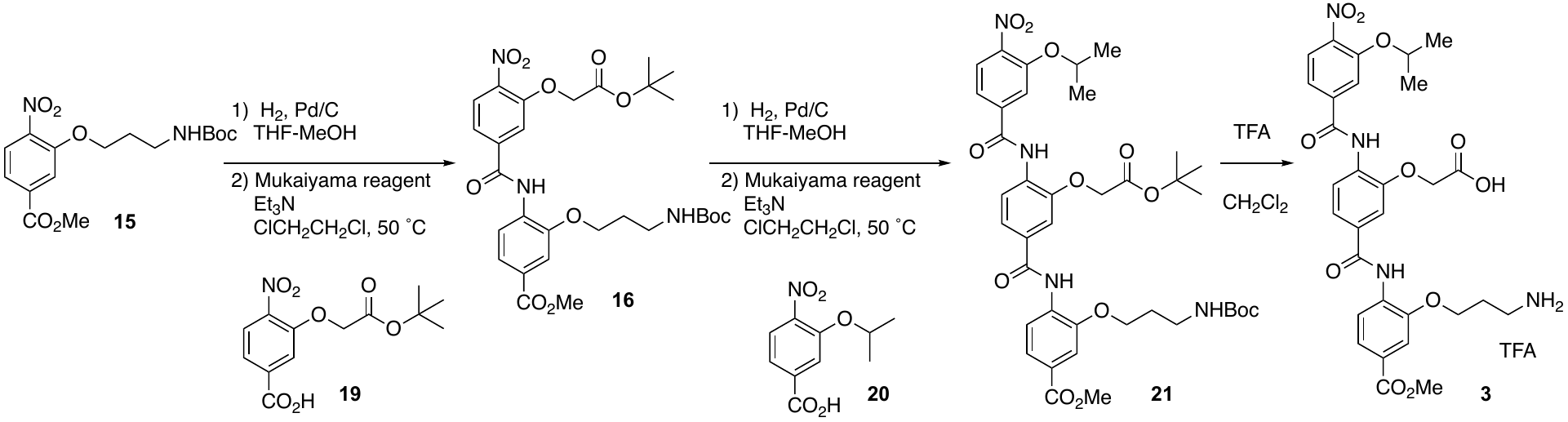

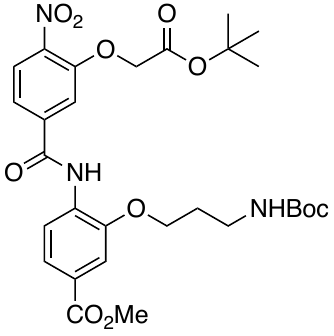
6 Methyl 4-(3-(2-(*tert*-butoxy)-2-oxoethoxy)-4-nitrobenzamido)-3-(3-((*tert*-butoxycarbonyl)amino) propoxy)benzoate (16)

According to general procedure (a), methyl 3-(3-((*tert*-butoxycarbonyl)amino)propoxy)-4-nitrobenzoate ^[4]^ **15** (1.00 g, 2.94 mmol) was reduced to the aniline which was then coupled with 3-(2-(*tert*-butoxy)-2-oxoethoxy)-4-nitrobenzoic acid **19**^[5]^ (1.05 g, 3.53 mmol). The residue was purified by flash column chromatography (3:7 to 3:2 EtOAc : hexane) to give *the title compound* **16** (1.21 g, 2.00 mmol, 68 % yield) as a yellow solid: δ_H_ (400 MHz, CDCl_3_) 9.05 (1H, br), 8.41 (1H, d, *J* 8.5), 7.95 (1H, d, *J* 8.2), 7.62-7.73 (3H, m), 7.53 (1H, s), 4.70 (2H, s), 4.55-4.65 (1H, m), 4.13 (2H, t, *J* 5.7), 3.85 (3H, s), 3.33-3.38 (2H, m), 1.90-2.03 (2H, m), 1.41 (9H, s), 1.29 (9H, s); δ_C_ (101 MHz, CDCl_3_) 166.5, 166.3, 163.0, 156.1, 151.4, 147.4, 141.9, 139.5, 131.5, 126.3, 126.0, 123.3, 119.9, 119.4, 114.7, 111.8, 83.2, 79.6, 66.6, 65.3, 52.1, 36.9, 29.5, 28.2, 28.0; IR 3429, 2979, 1749, 1702, 1588, 1130, 1054, 790; HRMS (ESI) calculated for C_29_H_36_N_3_O_11_ [(M-H)+]: 602.2344 found 602.2353.

7 Methyl 4-(3-(2-(*tert*-butoxy)-2-oxoethoxy)-4-(3-isopropoxy-4-nitrobenzamido)benzamido)-3-(3-((*tert*-butoxycarbonyl)amino)propoxy)benzoate (21)

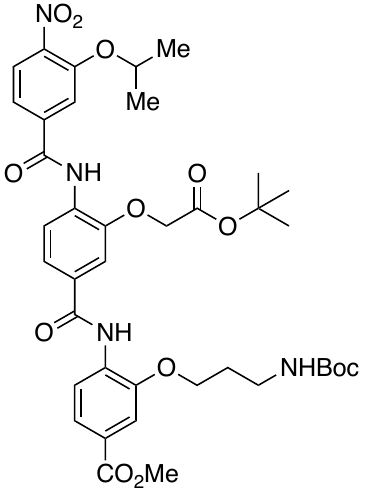
According to general procedure (a), methyl 4-(3-(2-(tert-butoxy)-2-oxoethoxy)-4-nitrobenzamido)-3-(3-((tert-butoxycarbonyl)amino)propoxy)benzoate **16** (470 mg, 0.776 mmol) was reduced to the aniline and then coupled with 3-isopropoxy-4-nitrobenzoic acid **20** ^[6]^ (210 mg, 0.931 mmol). The residue was purified by flash column chromatography (1:4 to 1:1 EtOAc : hexane) to give *the title compound* **21** (390 mg, 0.499 mmol, 64 % yield) as a yellow solid: δ_H_ (400 MHz, CDCl_3_) 9.53 (1H, s), 8.88 (1H, br), 8.58 (2H, d, *J* 8.4), 7.80 (1H, d, *J* 8.4), 7.74 (1H, s), 7.49-7.70 (5H, m), 4.79-4.93 (1H, m), 4.65 (2H, s), 4.59 (1H, br), 4.15 (2H, t, *J* 6.0), 3.85 (3H, s), 3.30-3.41 (1H, m), 2.01 (2H, d, *J* 6.1), 1.44 (9H, s), 1.37 (6H, d, *J* 6.1), 1.32 (9H, s); δ_C_ (101 MHz, CDCl_3_) 168.4, 166.6, 164.2, 163.3, 156.0, 151.3, 147.9, 147.1, 142.9, 139.0, 132.5, 132.0, 125.6, 125.3, 123.4, 121.4, 120.1, 119.4, 118.6, 115.2, 114.5, 111.6, 83.3, 79.5, 72.9, 68.4, 65.6, 52.1, 37.0, 29.6, 28.3, 28.0, 21.8; IR 3397, 2981, 1720, 1597, 1267, 748; HRMS (ESI) calculated for C_39_H_48_N_4_NaO_13_ [(M+Na)+]: 803.3110 found 803.3092.

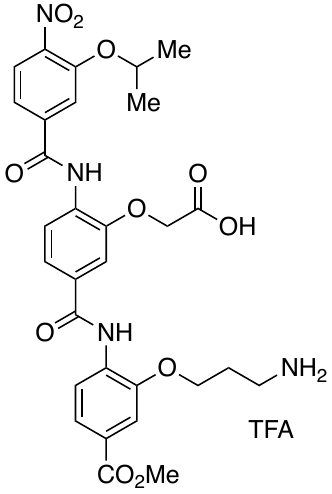
8 2-(5-((2-(3-Aminopropoxy)-4-(methoxycarbonyl)phenyl)carbamoyl)-2-(3-isopropoxy-4-nitrobenzamido)phenoxy)acetic acid 2,2,2-trifluoroacetic acid salt (3)

According to *general procedure (b)*, *t*-butyl compound **21** (390 mg, 0.499 mmol) was deprotected and purified by washing with dichloromethane/diethyl ether to give *the title compound* **3** (265 mg, 0.359 mmol, 72 % yield) as a yellow solid: δ_H_ (400 MHz DMSO-*d*_6_) 10.13 (1H, s), 9.63 (1H, s), 8.16 (1H, d, *J* 8.7), 8.05 (1H, d, *J* 8.2), 8.01 (1H, d, *J* 8.2), 7.87 (1H, s), 7.78 (2H, br), 7.59-7.73 (5H, m), 4.93-5.05 (1H, m), 4.90 (2H, s), 4.25 (2H, t, *J* 5.9), 3.89 (3H, s), 2.98-3.11 (2H, m), 2.04-2.16 (2H, m), 1.34-1.40 (6H, d, *J* 6.1); δ_C_ (101 MHz, DMSO-*d*_6_) 171.3, 166.3, 164.7, 164.1, 158.9, 150.3, 150.0, 149.8, 142.9, 139.4, 132.1, 131.6, 131.3, 126.5, 125.4, 123.4, 123.0, 122.9, 122.6, 121.8, 120.0, 115.6, 114.4, 112.7, 73.0, 67.6, 65.9, 52.6, 36.7, 27.2, 22.0; IR 2361, 1715, 1521, 1201, 870; HRMS (ESI) calculated for C_30_H_33_N_4_O_11_ [(M+H)^+^]: 625.2140 found 625.2116.

Results and Discussion

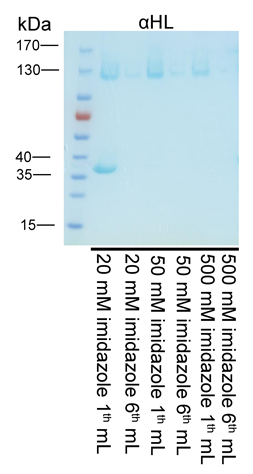

**Figure S1.** The purification of His-tagged WT αHL. Imidazole concentration gradient (20, 50 and 500 mM) was applied to elute αHL monomers and heptamers in the buffer of 50 mM Tris (pH 8.0), 500 mM NaCl and 3.8 mM DDM. Total 6 mL elution buffer was used to elute the supernatant from 100 mL culture for each imidazole concentration. The 1^st^ and 6^st^ mL elution buffer with different imidazole concentration was run on SDS-gel separately.

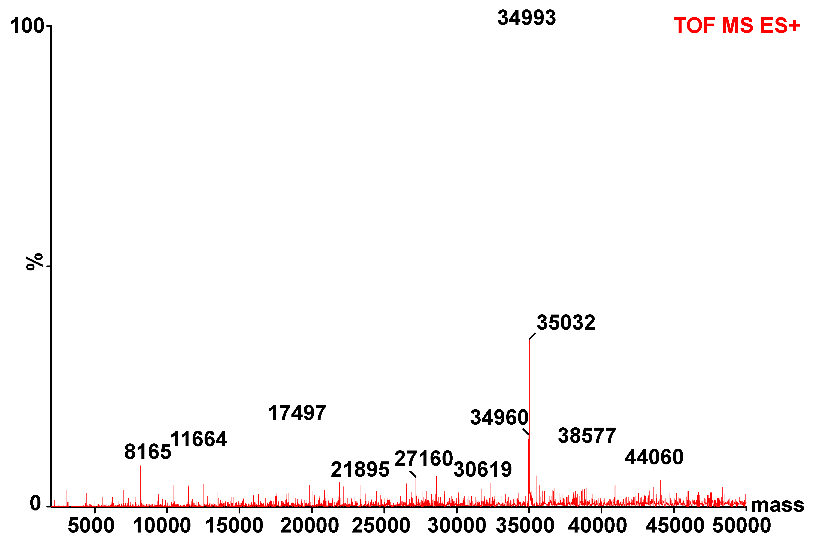

**Figure S2**. Deconvoluted mass spectrum of the purified WT αHL. LC-MS was conducted to confirm the molecule weight (MW) of αHL. The experimental molecule weight (MW) of αHL is 34993, compared to theoretical MW 34991 Da.

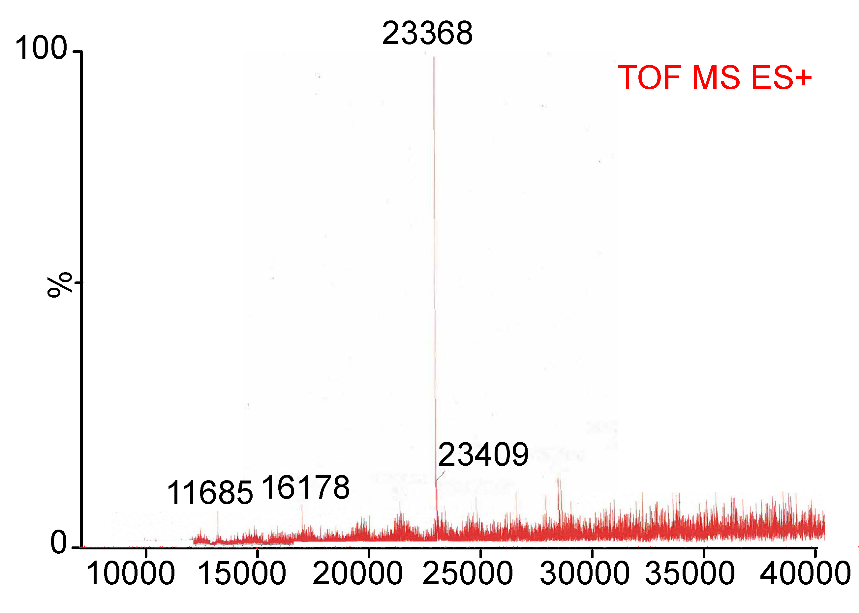

**Figure S3.** Deconvoluted mass spectrum of the purified nucleotide oligo (dC30) conjucted α-Syn αHL. LC-MS was conducted to confirm the molecule weight (MW).

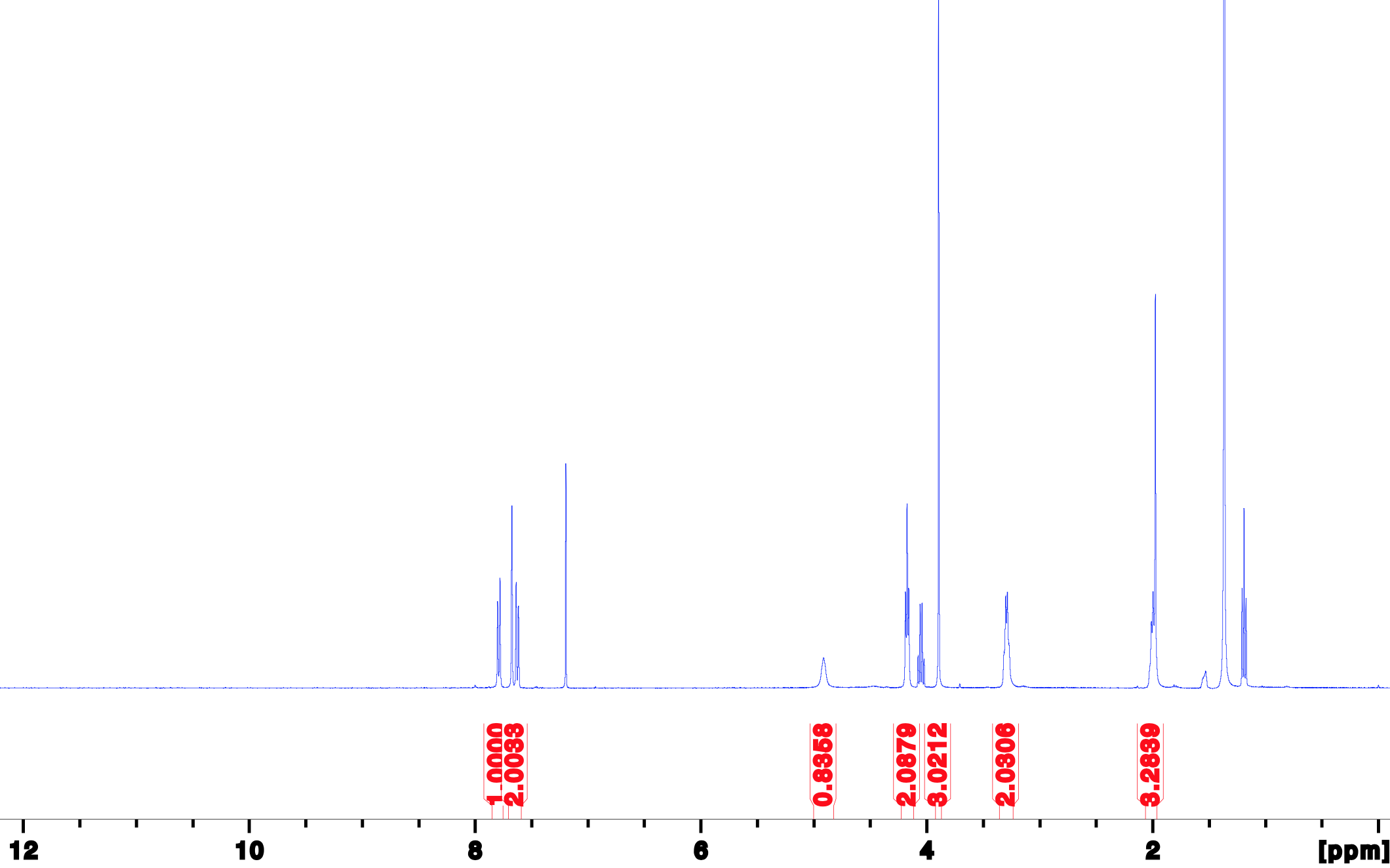
**^1^H, 400 MHz, CDCl_3_**

**Figure S4.** NMR spectrum of methyl 3-(3-((*tert*-butoxycarbonyl)amino)propoxy)-4-nitrobenzoate (15)

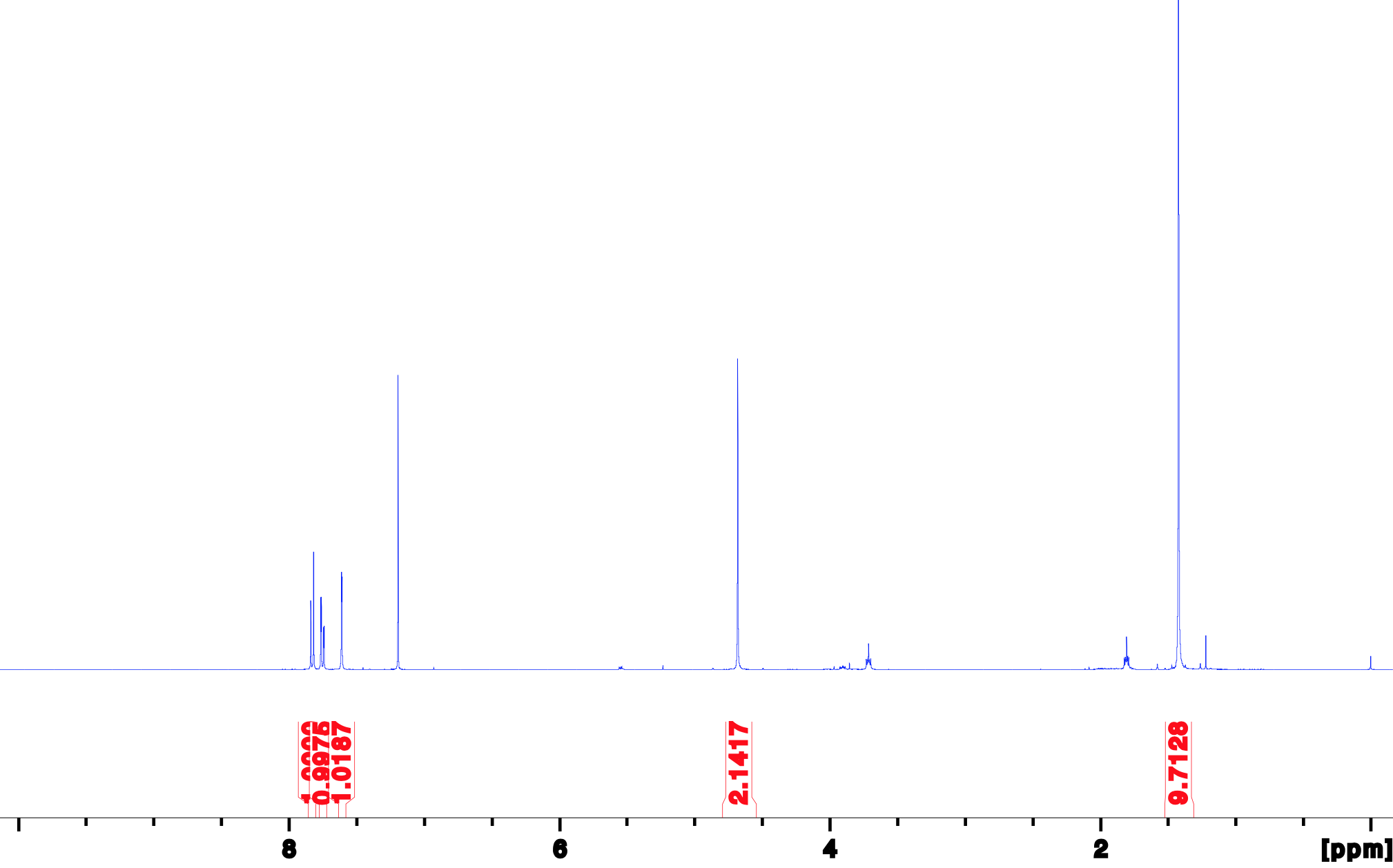
**^1^H, 400 MHz, CDCl_3_**

**Figure S5.** NMR spectrum of 3-(2-(*tert*-butoxy)-2-oxoethoxy)-4-nitrobenzoic acid (19)

**^1^H, 400 MHz, CDCl_3_**

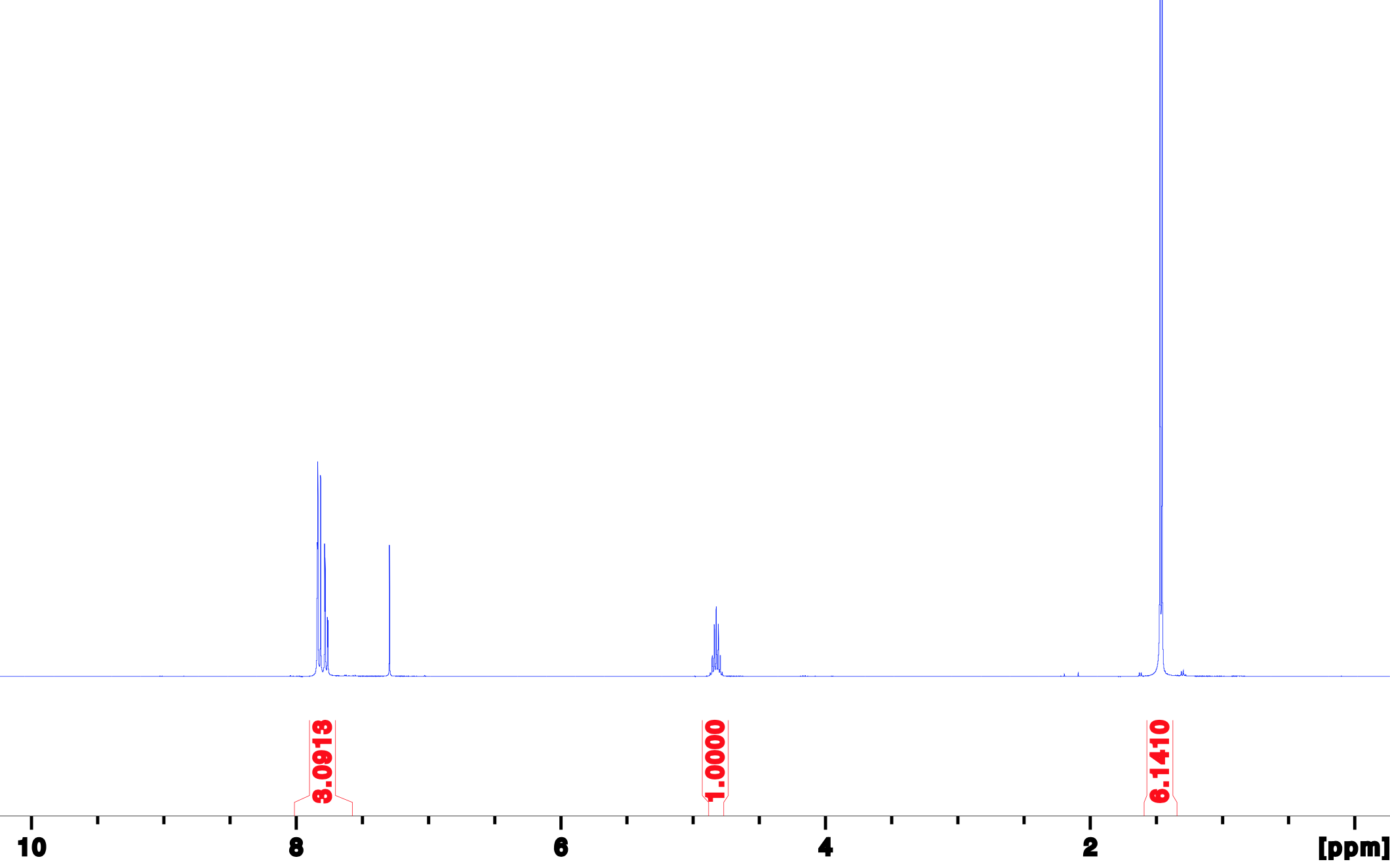
**Figure S6.** NMR spectrum of 3-isopropoxy-4-nitrobenzoic acid (20).

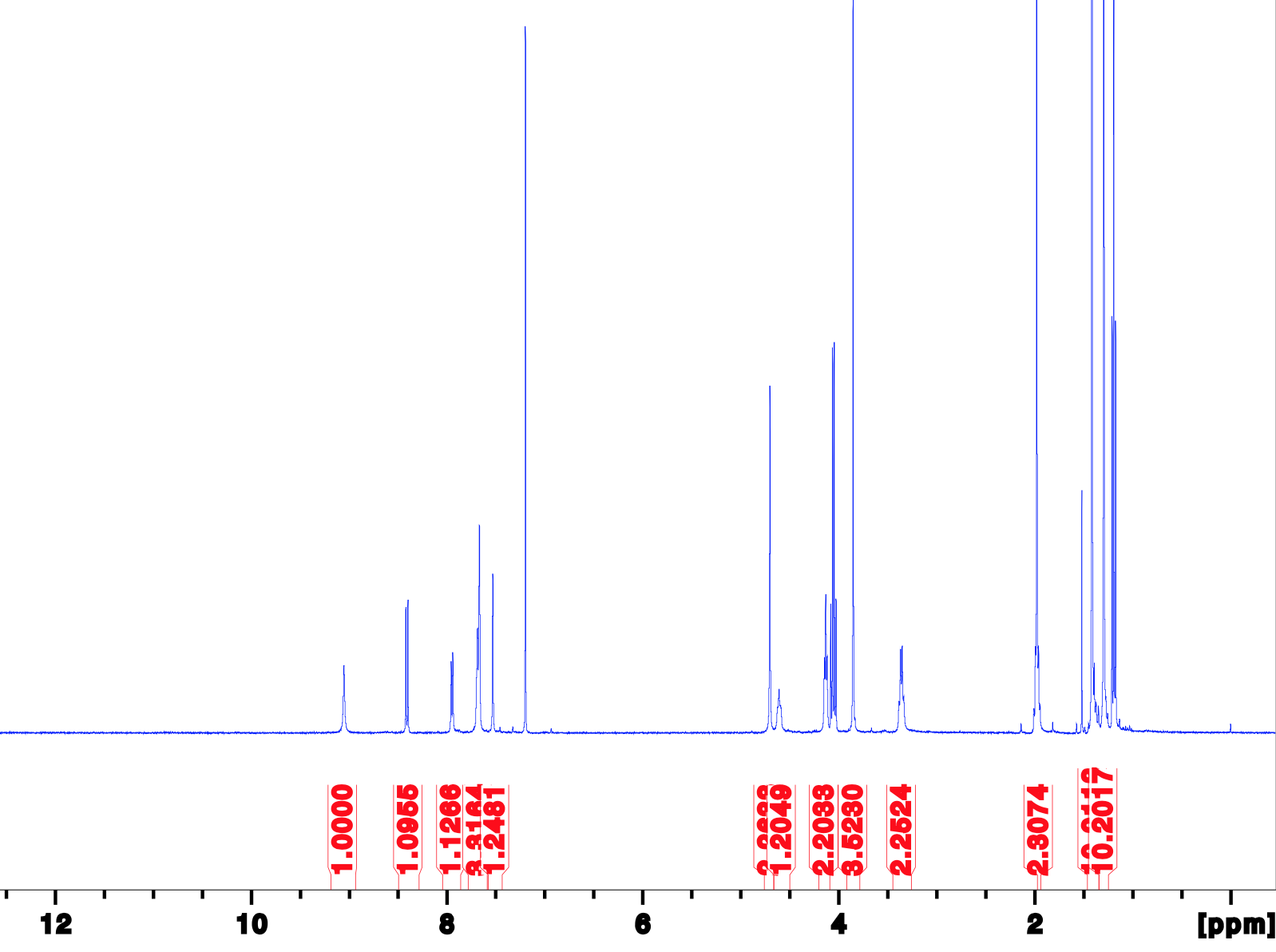
**^1^H, 400 MHz, CDCl_3_**

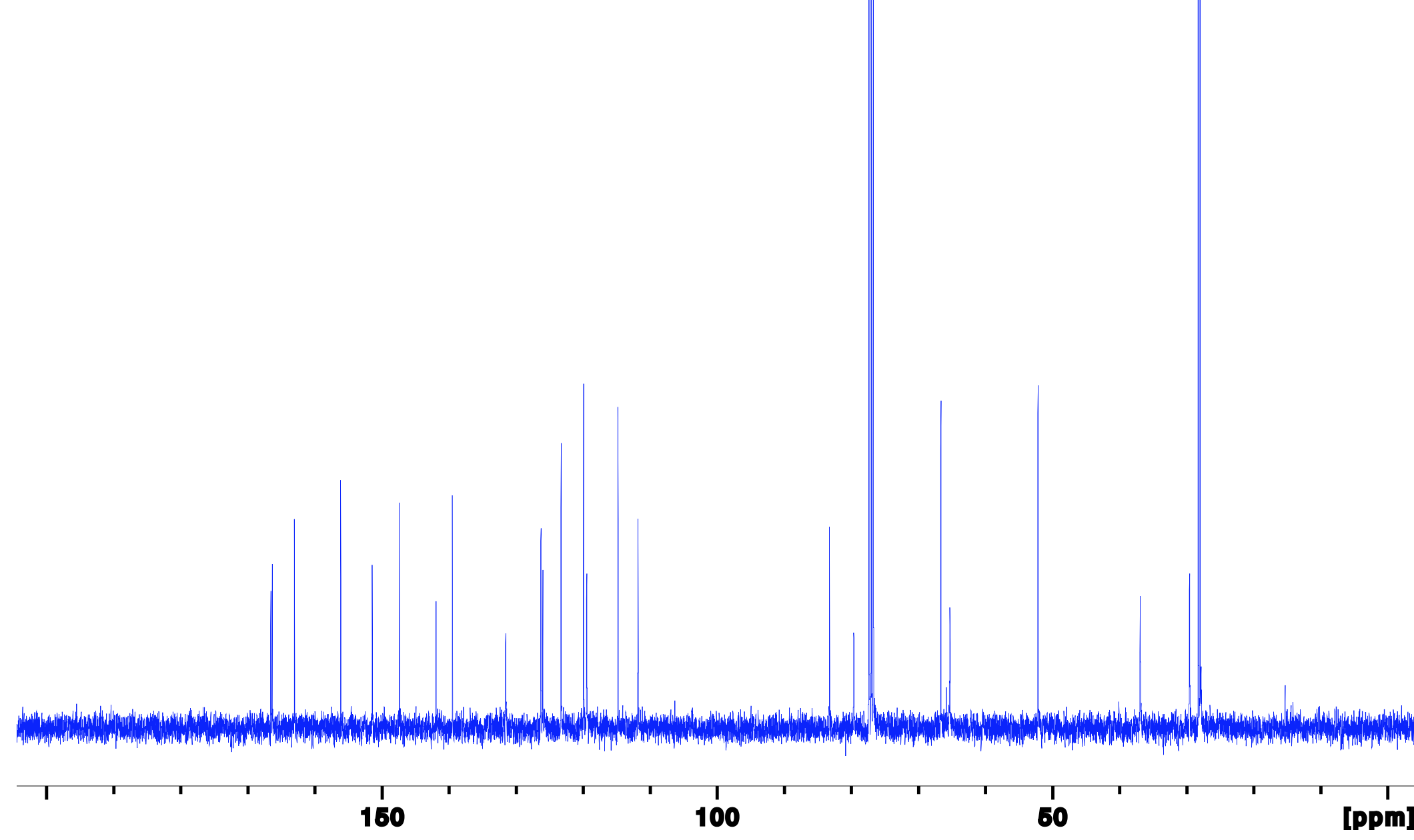
**^13^C, 101 MHz, CDCl_3_**

**Figure S7**. NMR spectrum (^1^H and ^13^C) of Methyl 4-(3-(2-(*tert*-butoxy)-2-oxoethoxy)-4-nitrobenzamido)-3-(3-((*tert*-butoxycarbonyl)amino)propoxy)benzoate (16)

**^1^H, 400 MHz, CDCl_3_**

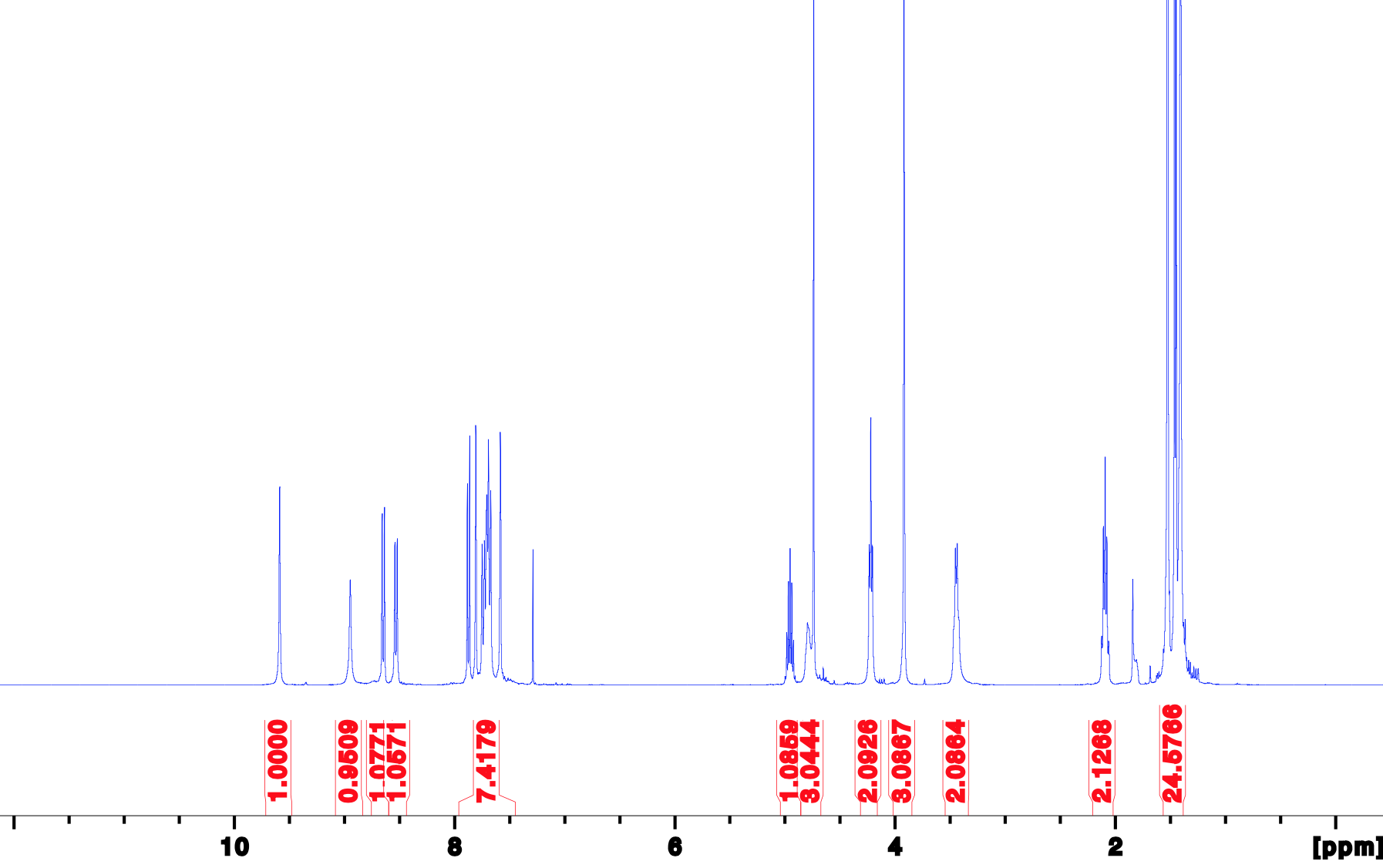
**^13^C, 101 MHz, CDCl_3_**

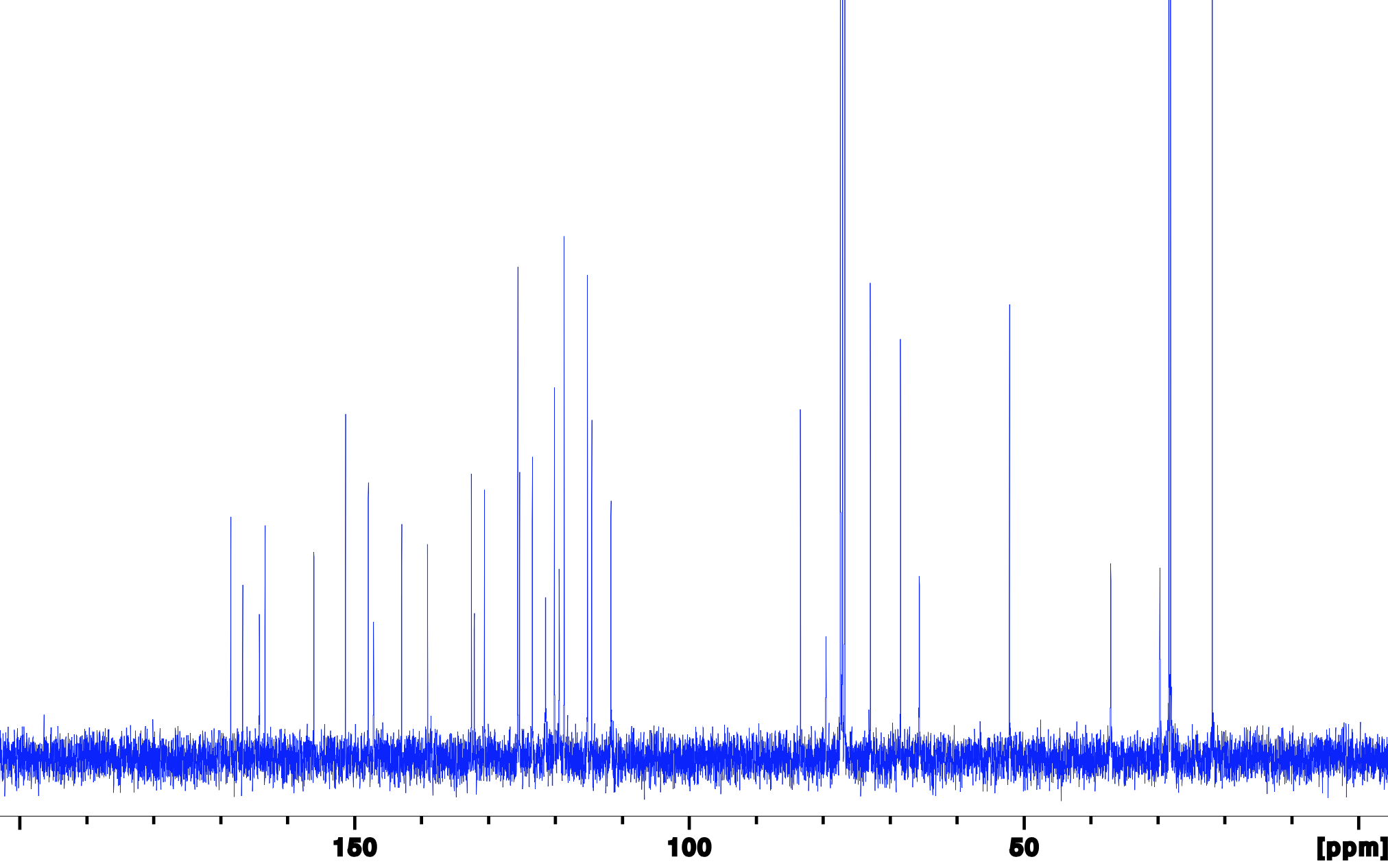
**Figure S8.** NMR spectrum (^1^H and ^13^C) of methyl 4-(3-(2-(*tert*-butoxy)-2-oxoethoxy)-4-(3-isopropoxy-4-nitrobenzamido)benzamido)-3-(3-((*tert*-butoxycarbonyl)amino)propoxy)be-zoate (21)

**^1^H, 400 MHz, d_6_-DMSO**

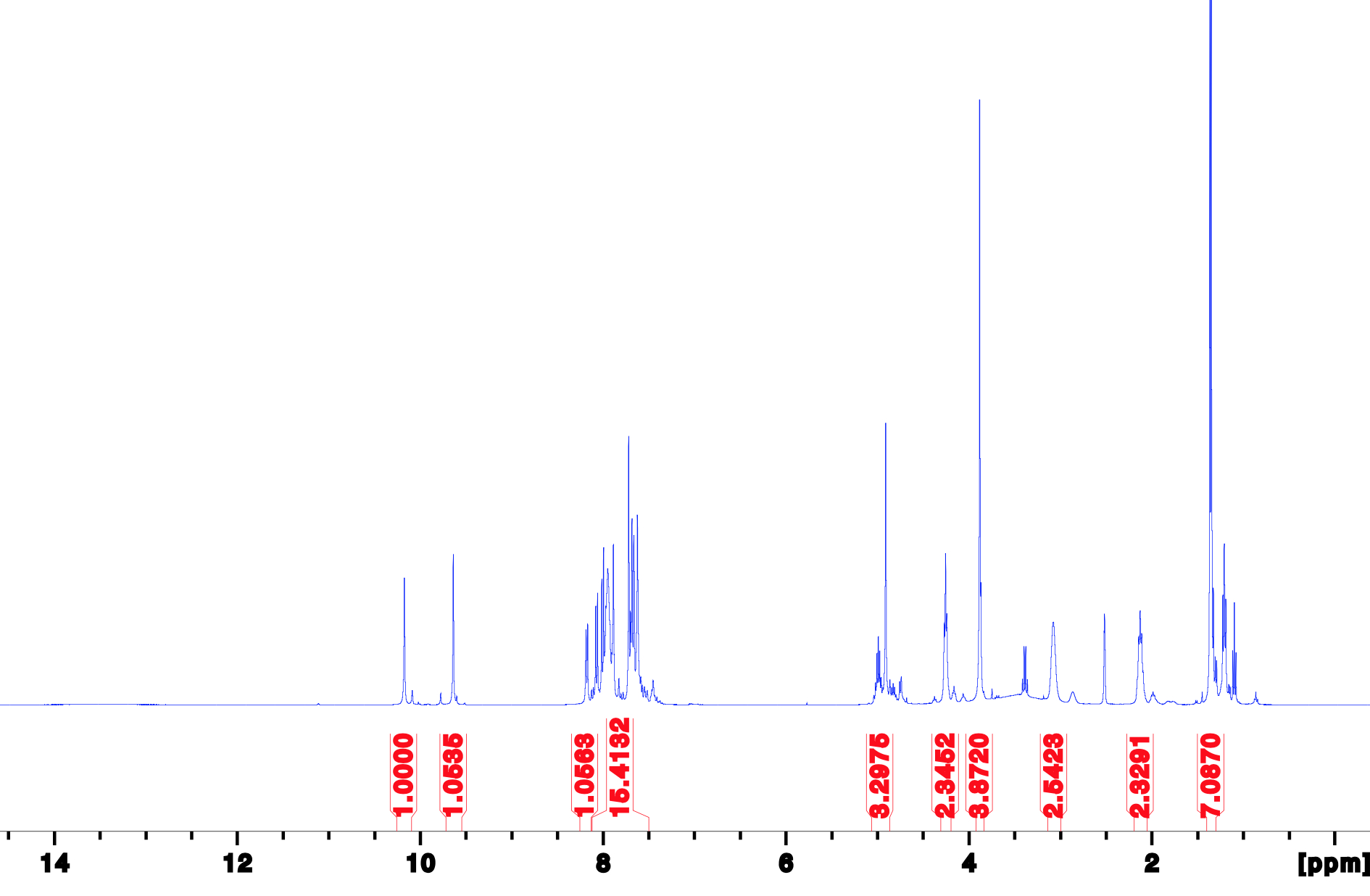
**^13^C, 101 MHz, d_6_-DMSO**

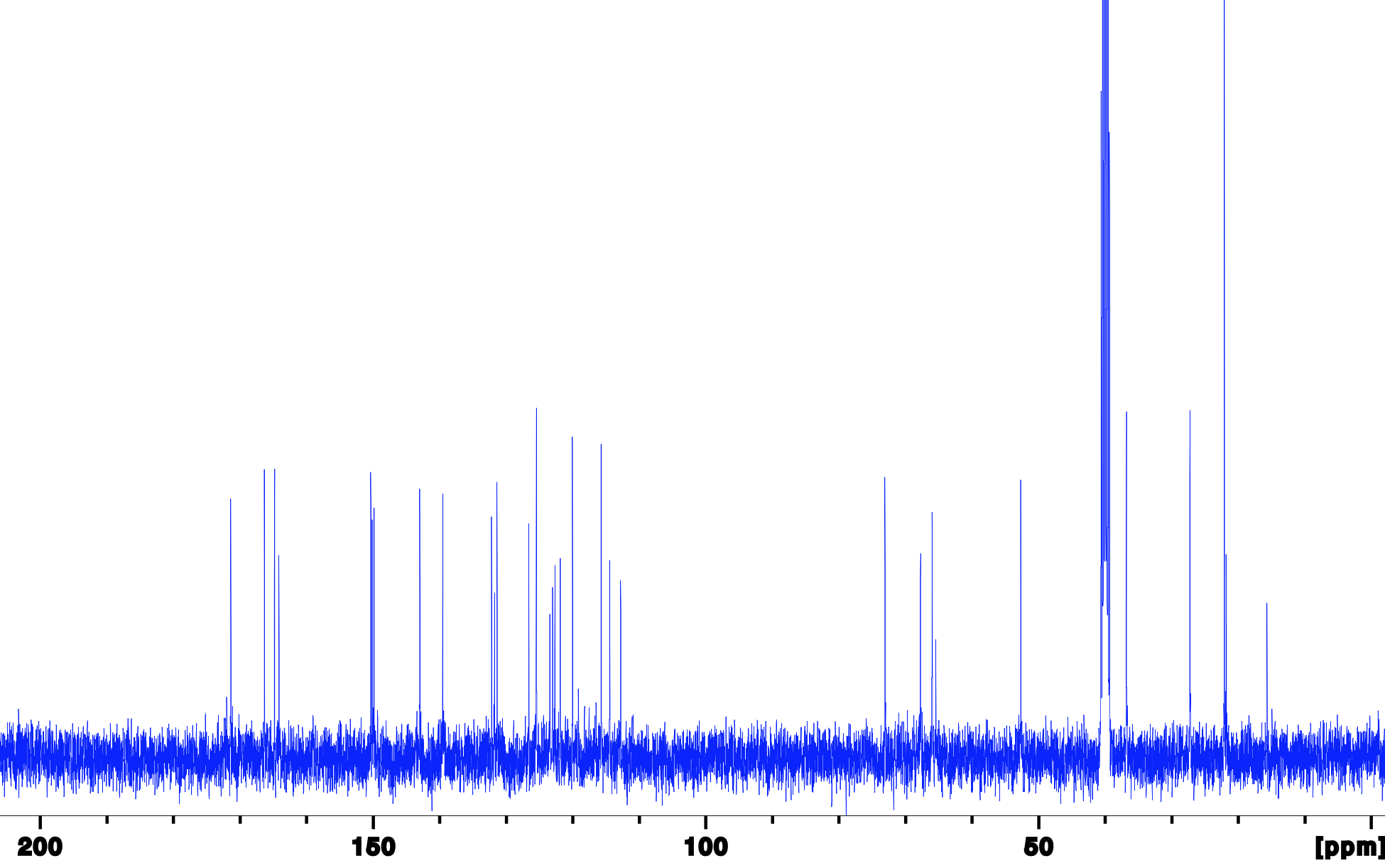

**Figure S9.** NMR spectra (^1^H and ^13^C) of 2-(5-((2-(3-aminopropoxy)-4-(methoxy-carbonyl)phenyl)carbamoyl)-2-(3-isopropoxy-4-nitrobenzamido)phenoxy)acetic acid 2,2,2-trifluoroacetic acid salt (3)
